# Sustainability of spatially distributed bacteria-phage systems

**DOI:** 10.1101/495671

**Authors:** Rasmus Skytte Eriksen, Namiko Mitarai, Kim Sneppen

## Abstract

Virulent phages can expose their bacterial hosts to devastating epidemics, in principle leading to a complete elimination of their hosts. Although experiments indeed confirm large reduction of susceptible bacteria, there are no reports of complete extinctions. We here address this phenomenon from the perspective of spatial organization of bacteria and how this can influence the final survival of them. By modeling the transient dynamics of bacteria and phages when they are introduced into an environment with finite resources, we quantify how time-delayed lysis, the spatial separation of initial bacterial positions, and the self-protection of bacteria growing in spherical colonies favor bacterial survival. This suggests that spatial structures on the millimeter and sub-millimeter scale plays an important role in maintaining microbial diversity.

**Author summary:** For virulent phage that invade a bacterial population, the mass-action kinetics predict extinction for a wide range of infection parameters. This is not found in experiments, where sensitive bacteria repeatedly are seen to survive the first epidemics of phage attack. To explain the transient survival of infected bacterial populations we develop a combination of local mass-action kinetics with lattice models. This model includes population dynamics, a latency time between phage infection and cell lysis, spatial separation with percolation of phages as well as colony level protection on the sub-millimeter scale. Our model is validated against recently published data on infected *Escherichia Coli* colonies.

## Introduction

Naturally occurring bacteria live in spatially structured habitats: in the soil [1, 2], in our guts [3, 4], and in food products [5]. The spatial heterogeneity is in part generated by the diversity of the microbial world [6], in part by clusters of food sources, and in part caused by the fact that bacterial division leaves the offspring close to their “mother” and thereby often form microcolonies [7–9]. The spatial heterogeneity may in turn amplify itself through propagation of host specific phages, if these percolate devastating infections through the parts of space with most homogeneous distribution of their hosts [10].

Traditionally, phage-bacteria ecosystems are modeled by generalized versions of the classical Lotka-Volterra equations [11–20]. In its simplest form, such mass-action equations predict sustained oscillations which becomes damped when one takes into account resource limitations. In contrast to the oscillating lynx-hare systems from macroscopic ecology [21], the microbial ecology experiments appear much more damped. However, with realistic parameters for phage infections [22] and realistic bacterial starting populations, the Lotka-Volterra type equations predict that an invading phage typically cause a collapse of the bacterial population to less than one bacterium, i.e. extinction. This is clearly not seen in microbial experiments. Rather, after an initial collapse to a measurable fraction of susceptible bacteria [14] they ultimately grow to a high density which matches the steady state prediction of the generalized Lotka-Volterra equations. These large initial oscillations can be mitigated by the inclusion variability in the adsorption rate of phages, e.g. by assuming two sub-species in a bacterial population, one with a high adsorption rate and one with a low adsorption rate [20]. However, this suggestion has the same issue as all mass-action models, namely that they ignore the spatial constraints in the physical system. Consequently, the models fail to capture that propagating phages tend to systematically deplete nearby hosts and thus may be unable to reach more distant hosts before the phages decay. In our paper, we consider spatial structures, both on the millimeter and sub-millimeter scale, as ways to moderate the virulence of the phages.

A typical method for introducing space in bacteria-phage modelling is by the use of cellular automata models [10, 23–25], but such models lack exponential growth of bacteria as only the bacteria adjacent to empty sites can replicate. By utilizing a combination of lattices, diffusion of nutrient and phages, and the shoving of bacteria, these models are used to study the interface between bacteria and phages. We propose a way to bridge the gap between the zero-dimensional mass action models and the spatial cellular automata models, and thereby incorporate both the spatial structures and the exponential growth of bacteria. With our model, we resolve much larger scale dynamics while retaining (some of) the sub-millimeter behavior of these models.

In particular, we are interested in systems where the bacteria form microcolonies, as in a semisolid medium or when bacteria stick together due to the formation of biofilm. In these scenarios the sub-millimeter structure of the colonies requires additional modifications to the traditional Lotka-Volterra models which we will describe below.

We partition space into a three dimensional lattice, which allows for spatial variation in the densities of phage and bacteria (see Fig. 1A). Consequently, the fallout of events needs to propagate before interacting with distant areas. Each box in the lattice is well-mixed, meaning that bacterial colonies within each box have the same composition, but this composition is typically different from colonies in other boxes. Due to their small size, phages and nutrients diffuse readily around in the system, so we include diffusion to couple the dynamics in one box with its neighboring boxes (see Fig. 1B).

**Fig 1.**
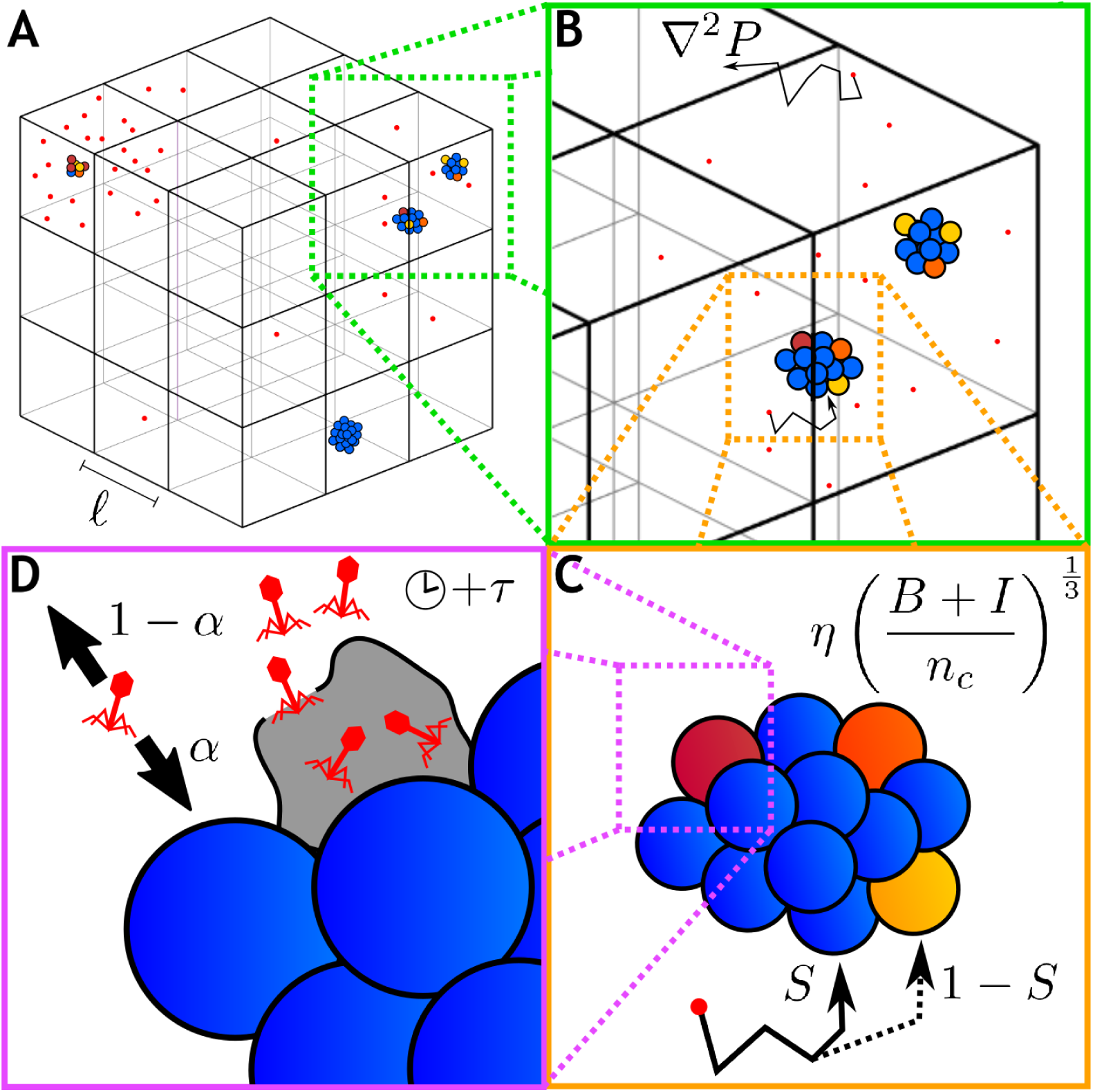
Schematic of new concepts. (A) Spatial variation is included by the introduction of a three dimensional lattice where the dynamics of the bacteria-phage interaction plays out largely independently in the separate boxes. (B) Within each box we imagine bacteria growing as small colonies (colored circles) with the phages (red dots) and nutrient (not shown) interacting freely within the box. All colonies within a box are assumed to be identical but they may differ from colonies in other boxes. Diffusion couples the phages and nutrients in one box with neighboring boxes. (C) Adsorption to a single large target has different properties compared to adsorption to several small targets and consequently the phage adsorption term is proportional to the bacterial density to the power of 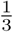. Furthermore, the sub-millimeter structure of the colony also modifies the probability for the phage to hit uninfected cells (parameterized by the function *S*). (D) Cell lysis occurs after a latency time of *τ* upon which phages are released from the lysed cell. Of these phages, a fraction 1 − *α* fully escape the colony and the remaining fraction, *α*, immediately find new targets to infect.

When bacteria form colonies the dynamics of phage interaction changes in several ways:

1. When phages adsorb to a single large target they do so with a rate which is proportional to the radius of the target as derived by Smoluchowski [26, 27], thus proportional to the volume to exponent 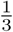. This reduces the adsorption rates to much smaller than in unstructured situations (see Fig. 1C).
2. Infected bacteria typically lie on the surface of colonies rather than being uniformly distributed throughout the colony [9, 23]. This means that the probability of a phage hitting an uninfected cell is not just proportional to the relative proportions of uninfected to total cells, but is reduced by the shielding function *S*, which we discuss in the methods section (see Fig. 1C).
3. When progeny phages are released from a colony, the phages will have many new targets readily available locally. We therefore include a readsorption effect where a fraction of the released colony immediately reinfect new bacteria while the remaining phages escape fully from the colony (see Fig. 1D).

Our model combines all of these aspects to describe phages attacking confined bacteria.

One of the useful model systems of spatially structured microbial population is found in the human gut. For example, during fecal microbiota transplantation, new bacteria and phages are introduced to the patients gut in an attempt to restore diversity to the microbiome. The subsequent fate of a phage epidemic is played out in the spatially constrained environment consisting of bacteria that divide in relatively fixed positions, while potentially depleting the local nutrient as the population grows to the carrying capacity of the niche. The success of the treatment may be influenced by the phage epidemic as their ability to control bacterial biomass can alter the microbiome [28].

## Materials and methods

A typical set of mass-action equations which describes the bacterial density *B*, the phage density *P*, and the nutrient *n* could take the form [13, 14, 20, 29]:

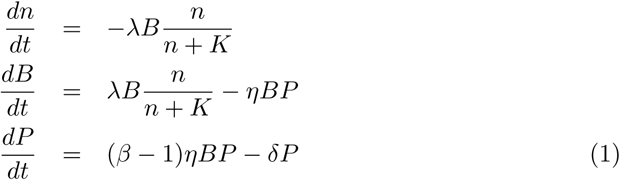

where the nutrient *n* is measured in units of bacteria it can be converted to (corresponding to a yield of 1), the *λ* = 2 *h*^−1^ is the maximal bacterial growth rate, *η* = 10^−8^ *µm*^3^/*h* is the adsorption rate for phages, *β* = 100 is the phage burst size, and *δ* = 0.1 *h*^−1^ is the phage decay rate [22, 30]. *K* = *n*(0 *h*)/5 is the Monod constant [31] which determines at which concentration of nutrients the growth rate is halved. The initial nutrient level is set to *n*(0 *h*) = 10^9^ /*ml*, meaning there is sufficient nutrient to produce 10^9^ bacteria within a milliliter of medium.

These equations are modified from standard Lotka-Volterra equations by the depletion of food, and the associated parameterization of growth by the Monod equation. The above equations are the simplest way to model the development of a bacteria-phage system in a fixed batch culture (e.g. as used for temperate phage dynamics in [29]).

Even before considering the effects of space on bacteria-phage interaction, the model noticeably ignores the latency time between phage infection and cell lysis. This is easy to include, either through a time delay equation [32] or by introducing some intermediate infected states, *I*_*k*_, between infection and lysis [33, 34]. We choose the latter method. In addition to the latency, we also include millimeter-scale spatial effects (density variation and diffusion) and sub-millimeter scale spatial effects related to bacterial colonies. Taken together, we present our full model for phage-bacteria interaction in a spatial context.

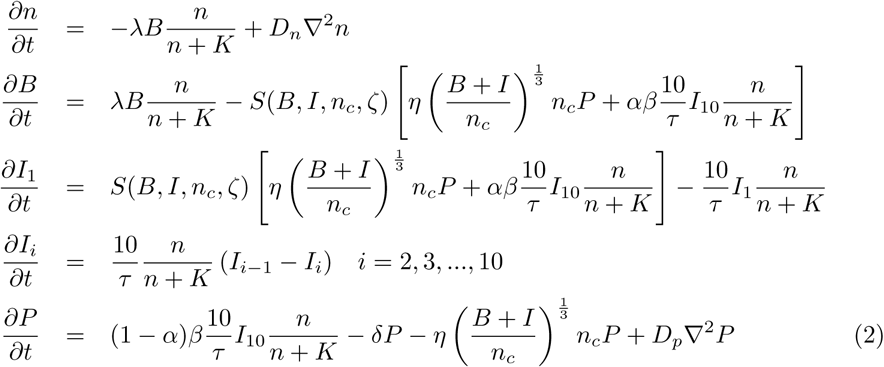

The new concepts included in this model is shown schematically in Fig. 1 as guide to the reader. The populations (*B,P, I*_1_, …, *I*_10_) and the nutrient *n* are now functions of space (*x, y, z*) as well as time, but we keep the parameters constant throughout. For readability we don’t explicitly include the spatial dependency in the notation. In the model, *D*_*n*_ and *D*_*p*_ are the diffusion constants for the nutrient and phages respectively. Here *D*_*n*_ ∼ 10^6^*µm*^2^/*h* [35] is much larger than *D*_*p*_ ∼ 10^4^*µm*^2^/*h* [36, 37].

We investigate scenarios where the bacteria are constricted in the medium and remain effectively immotile. This situation is for instance encountered when performing experiments in soft agar [9], but there are several other systems where the bacteria are constrained (such as soil, biofilm, solid foods). This constriction in space means that each initial bacteria grow to a colony of size *(B* + *I*)/*n*_*c*_, where *n*_*c*_(*x, y, z, t*) = *B*(*x, y, z*, 0 *h*) is the density of the initial bacteria and thus the density of colonies (see Fig. 1B), and *I* is the density of infected cells.

When bacteria grow in colonies as opposed to growing separately, the dynamics of phage adsorption changes significantly. When growing as a colony, the bacteria presents a smaller surface to the outside compared to growing individually. Therefore, when a phage is adsorbed to one of the *n*_*c*_ colonies, it does so with a rate that is proportional to the colony radius [26, 27] (hence the 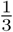 exponent, see appendix for more details).

When some bacteria in the colony are already infected, then only a fraction *S* of the infections leads to new infected states, since the accumulated infected cells exists on the surface of the colony and partially shields the uninfected cell from the phage attack [9]. This protection is defined by the shielding function *S*, which we will explain further below.

The model in Eq. (2) also includes the latency time of the infection cycle, which is obtained with the use of 10 intermediate infected states 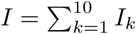 between infection and final release of phages [34]. The use of intermediate stages leads to the latency times being drawn from gamma distribution with an average latency time *τ* = 0.5 *h* [22] and a variation of *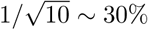* between separate lysis events. The latency between infection and lysis is furthermore scaled by a Monod-like term, meaning that as the nutrient is depleted, the latency time increases with bacterial generation time and thereby the phage invasion is slowed. When lysis occurs, the released phage has a high probability of reinfecting the colony, which is represented by the parameter *α* = 0.5. A fraction (1 − *α*) of the phage progeny escape from the colony and becomes free phages *P*, while a fraction *α* readsorbs to bacteria in the colony. The functional form of the readsorption as well as the value of *α* is somewhat arbitrary, since we do not have a good model for what it means to “escape” a colony, nor do we have a model for how many phages should readsorb within a reasonable timescale. Instead, we use *α* as a simple way to penalize colony growth. In the appendix, we investigate several values of *α* and find that the survival of the bacteria is largely unchanging for *α* values between 0.5 and 0.95.

Returning to the choice of shielding function *S*, we investigated various shielding functions (see appendix for details), and found an approximate shielding function of the form:

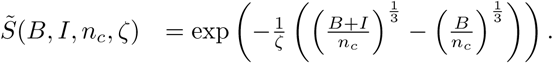

This shielding function gives the probability that a diffusing phage passes through the absorbing surface of thickness 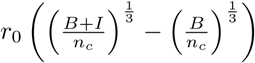, where *r*_0_ is the radius of a single bacterium. The parameter *ζ* is the typical distance that a phage can penetrate before being adsorbed (in units of *r*_0_). In the Appendix, we estimate *ζ* as a function of the phage adsorption coefficient via detailed simulations of colonies consisting of spherical bacteria, and find that for the diffusion limited case *ζ* is roughly 1.

Note that to match in the shielding in the small colony limit where 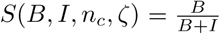, we must force an upper limit on the shielding:

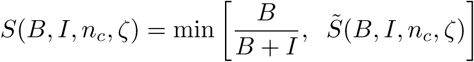

In our implementation of the model, we include the spatial variation by simulating a one cubic centimeter system where space is sub-divided onto a 3-dimensional lattice consisting of boxes of length *ℓ* = 0.2 *mm*. This approach allows for different numbers of bacteria and phages in each of the 1.25 · 10^5^ boxes in the lattice. Note that in order to represent the behaviour of a larger system we use periodic boundary conditions.

Inside each box in the lattice we simulate our model stochastically by treating each term in Eq. (2) as rates corresponding to events. At each time step, we compute the probability for every particle to undergo each event, and draw the number of occurrences from a Poisson distribution (see appendix for details). Every box is simulated independently, modulated by diffusion of food and phages between the boxes. The diffusion of phages is done by random walk of appropriate fractions of the phage population to neighboring lattice boxes. Nutrient diffusion is computed by using a discrete Laplace operator (using the “forward time central space” scheme).

The problem contains two different timescales, that of bacteria and phages versus that of nutrient diffusion, which is the fastest timescale. In order to solve nutrient diffusion accurately we require that 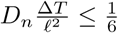, which we obtain for time steps of size Δ*T* = 2 · 10^−3^ *h* ∼ 7 *s*.

In Fig. 2, we show an example of a single simulation. Here the spatial heterogeneity is evident both in the separation of bacteria colonies in the lattice (blue spheres) but also in the distribution of phages (red points): the phages form small clouds as they diffuse away from areas where the invasion has taken hold.

**Fig 2.**
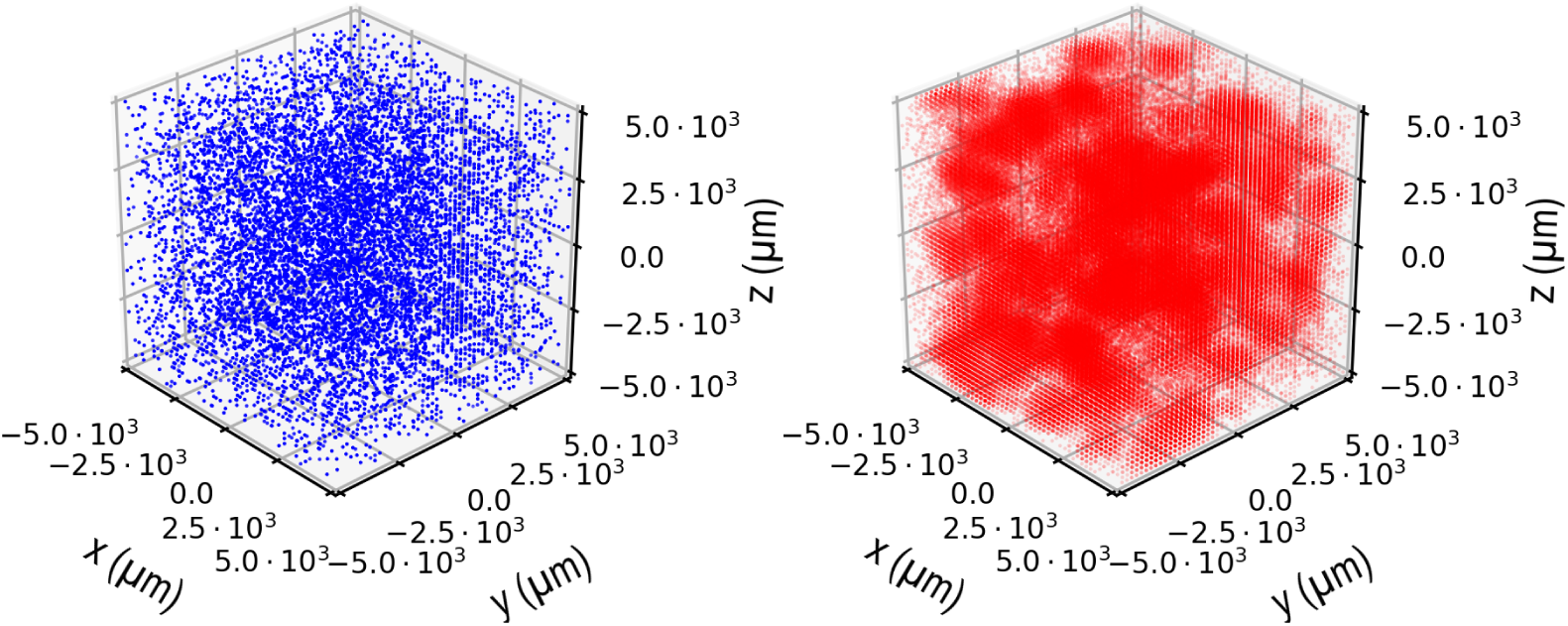
Snapshot of simulation. The spatial distribution of bacteria (blue) and phages (red) at time *T* = 7 *h* starting from an initial *B* = 10^4^ bacteria per ml and *P* = 10^5^ phages per ml. The opacity of each point is proportional to log 10 of the population in that point.

The model we present in Eq. (2) has two limiting cases which we also simulate as part of investigating how space and time-delayed lysis effect the dynamics of the interaction.

In the first limiting case, we consider a situation where all of the sub-millimeter colony-level protection mechanisms are ignored, i.e. we set 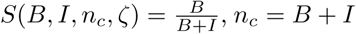, and *α* = 0. In this limit the term: 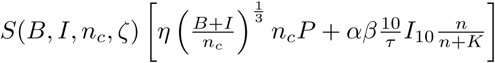 reduces to well-mixed interaction term *ηBP*. Under these conditions, we now model how delayed lysis and spatial heterogeneity in the population densities moderate the virulence of the phages (see e.g. Fig. 4C). This limit corresponds to the situations where the bacteria are not immobilized and thus do not form colonies, e.g. as in liquid medium. The spatial heterogeneity then comes from the difference of diffusion rates, where bacteria, due to their large size, have diffusion rates between ∼ 3 · 10^2^ *µm*^2^/*h* and ∼ 7 · 10^2^ *µm*^2^/*h* [38] (assuming no chemotaxis). Since our lattice is resolved at a resolution of ℓ = 200 *µm*, we can approximate the bacteria as being effectively immotile since they diffuse on average ∼ 25 *µm* per hour and thus would be unlikely to travel much further than to the boxes immediately adjacent.

We can take this limit further by setting the lattice size *ℓ* to be equal to the system size, thereby reducing our lattice to a single box which removes all spatial heterogeneity while retaining the mechanism of delayed lysis (see e.g. Fig. 4B). A typical example of this final limit would be bacteria growing in a liquid culture in a shaker incubator.

## Results

The model is simulated with four different levels of spatial detail. Each level shows the effects of the changes from the initial mass-action equations we presented in Eq. (1), to the full spatiotemporal model that also include protection due to colony formation.

Examples of simulations are illustrated in Fig. 3, where one can follow the trajectory of the system as described by the different levels of details: In (A) we solve Eq. (1), and observe how the phage quickly overwhelms the bacteria leading to the extinction of the bacteria. In (B) we solve Eq. (2) without the addition of space (one box with *ℓ* = 1 *cm*), and we also remove the effect of colony level protection by setting *n*_*c*_ = *B* + *I*, 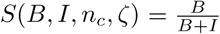 and *α* = 0. This leaves us with a mass-action model like Eq. (1) but which includes the latency between phage infection and cell lysis. The latency of the lysis cycle delays the growth of phages, thereby leaving more time for bacteria to grow. As a result, the bacteria reach a higher density and subsequently is exposed to a more dramatic collapse when phage population catches up. Thus the inclusion of the appropriate time delays predicts more unstable behavior, and is accordingly less in agreement with typical experiments than the simpler equation used in panel (A). Panel (C) and (D) explores the two levels of spatial detail, both with the appropriate time delay due to latency. Panel (C) only introduce spatial variation and keep the restraints 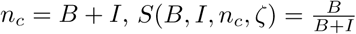, and *α* = 0. The added heterogeneity of space further delays the extinction of the bacteria as different regions of space experience different phage densities. Panel (D) use the full model that includes colony level protection. One sees dramatic improvement in stability, with bacteria surviving to reach a steady state where normal phages cannot propagate. Notice that we do not model large plaque former like phages T7 here, as we assume that phage latency diverge with same Monod factor as the bacterial generation time (Eq. (2)). In fact, when including protection due to colony formation, the bacteria can only be eliminated when the initial phage load is high enough to eliminate bacteria before they form sufficiently large colonies.

**Fig 3.**
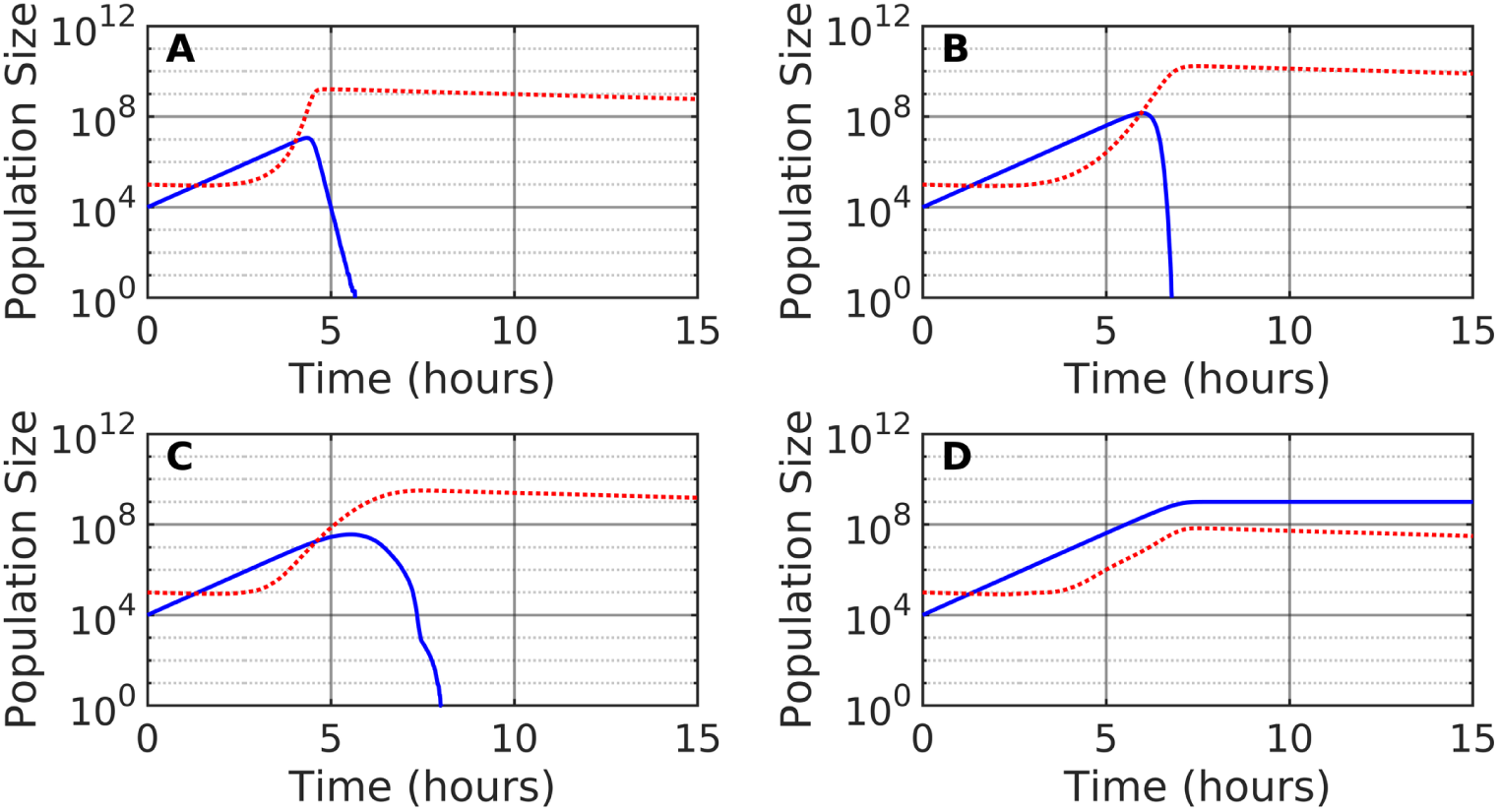
Dynamics of bacteria and phages. The solid blue lines indicate the bacterial population, while the dotted red lines indicate the phage population. (A) Simple Lotka-Volterra. (B) Lotka-Volterra with a time-delay. (C) Lotka-Volterra model with time-delay and space. (D) Lotka-Volterra model with time-delay, space, and colony level protection. In all cases we start with the initial condition *B* = 10^4^ bacteria per ml and *P* = 10^5^ phages per ml, and assume a max production of *n*(*t* = 0) = 10^9^ bacteria per ml.

Fig. 4 explores bacterial survival as function of initial density of phages and bacteria in the four levels of modeling detail. In each case, the simulation is initiated with bacterial and phage populations between 1 and 10^9^ per ml. The spatial simulations consider 1 *cm*^3^ = 1 *ml* coarse grained into boxes of dimension 0.2 *mm*, and a bacterial population of 10^5^ per ml accordingly correspond to an average population of one bacteria per box. In each panel one can see a dark blue region that represents conditions where bacterial populations are entirely eliminated. For each model variant this “dead” region expands as simulation time is increased (going from left to right shows to the bacterial population at 5h, 10h, and finally 15h). After 15h the nutrient is depleted in most cases. This is highlighted by the red line, marking conditions where the average nutrient density is equal to the Monod growth constant: 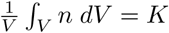. Above this red line the accumulated number of bacterial divisions approaches the carrying capacity, and the system remains largely unchanging.

**Fig 4.**
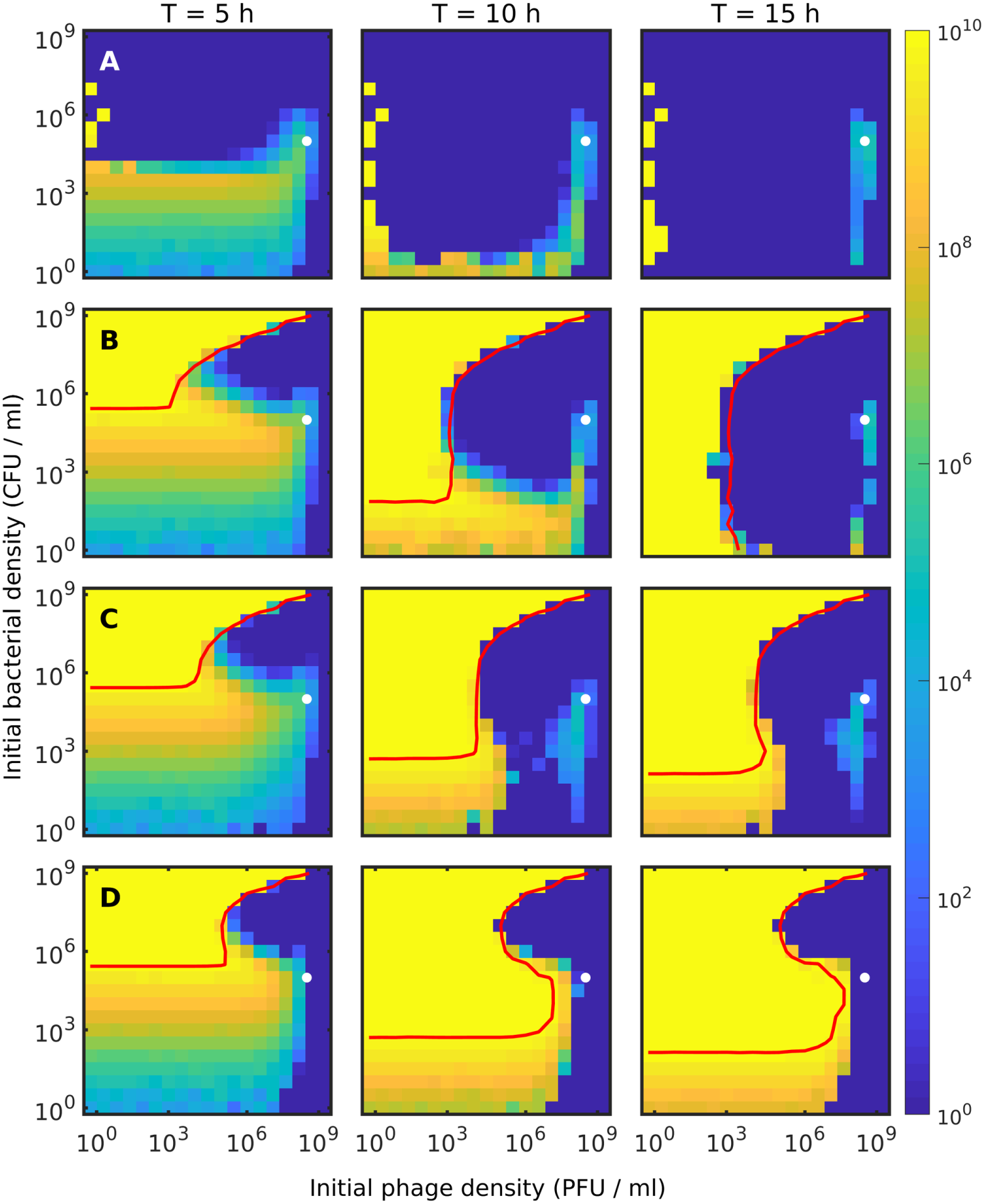
Phase space plots of the surviving bacteria. The color shows the average bacterial population density at the given time as a function of initial phage density and initial bacterial density (averaged over the whole system). The red lines indicate the nutrient contours 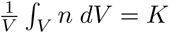, where the growth rate is halved. The white dots indicate where *P* = 2 · 10^8^ / *ml* and *B* = 10^5^ / *ml*. Each column show the dynamics at *T* = 5 *h, T* = 10 *h*, and *T* = 15 *h* respectively, while the rows indicate different models: A) Simple Lotka-Volterra. B) Lotka-Volterra with a time-delay. C) Lotka-Volterra model with time-delay and space. D) Lotka-Volterra model with time-delay, space, and colony level protection.

Fig. 4 (A) show that in the traditional mass-action models, the fate of the bacterial populations is similar across a range of initial conditions where the phage eradicate the bacteria. In some cases, we observe transient survival where bacterial growth balances death due to phages. In quantitative terms, the bacteria population is marginally sustained when the rate of phage adsorbing to bacteria is approximately equal to the rate of bacteria division, *λB* ∼ *ηBP* ⇒ *P* ≈ *λ/η* = 2 · 10^8^ / *ml*. Further, the phage population cannot increase when the rate of phage produced by lysis is less than the decay rate of the phage.

At the point where phage adsorption is balanced by the bacterial growth rate (*P* ≈ 2 · 10^8^ / *ml*), the population of infected bacteria is approximately *I* ≈ *λBτ*, which means that the phage production rate is approximately 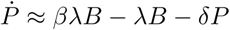. For our parameter values, this is zero when *B* ≈ 10^5^ / *ml*. When initial B is larger than 10^5^ / *ml*, the phage population can grow from the beginning, whereas lower bacterial density means that phages cannot grow before bacterial population has increased. Around these critical densities, the phage does not grow sufficiently fast to repress the bacterial population within 15 h.

Fig. 4 (A) Shows that the initial Lotka-Volterra dynamics only predict long term survival, when the phage density is extremely low. Here the bacteria can reach carrying capacity and thus survive indefinitely, due to the phages decaying before adsorbing to the bacteria.

Fig. 4 (B) demonstrate that the latency time of phage infection enables the bacterial population to survive in a much larger region of initial conditions, compared to panel (A). This survival arises because the latency allows the bacterial population to deplete the nutrient before the phage population grows large enough to exterminate the bacteria. This observation gives an addendum to our observation based on Fig. 3 (B): while latency leads to larger crashes of the bacterial population in some cases, it leads to long term survival in other cases.

Fig. 4 panels (C) and (D) demonstrate the moderating effects of space, where the initial conditions for long term survival for bacteria is greatly extended. The combination of sub-critical percolation at moderate levels of initial bacterial density (C), and the buildup of sizable colonies (D) tend to provide for robust bacterial survival also in the region around 10^4^ − 10^8^ initial phage per ml.

To test the model, we modify the setup to mimic the conditions used in the experiment performed in ref. [9]. There a number of *Escherichia coli* was suspended in a soft agar and plated onto petri dishes. After a time where each bacterial had grown to some small colony, the plates were sprayed with the virulent phage *P* 1_*vir*_, and the number of visible bacterial colonies were counted 16 hours later. In the experiment the formed bacterial colonies were about 1 *cm* apart, embedded in a soft agar of height ≈ 400 *µm*. These conditions is replicated by simulating a single bacterium inside a box of dimensions 10^4^ *µm* × 10^4^ *µm* × 400 *µm*. The phages are spawned in the XY plane at *Z* = 400 *µm*, and we change the boundary condition in the Z axis from periodic to reflective. In addition, we increased the spatial resolution by setting *ℓ* = 0.1 *mm* in order to better resolve the Z axis. The increased resolution requires the nutrient to be solved on a time scale of Δ*T* = 5 · 10^−4^ *h*. The shorter time scale means that computation time increases significantly. To remedy this situation, we exploit the fact that phages move on average 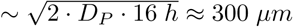 during the experiment. We therefore only simulate the phages in an area of 10^3^ *µm* × 10^3^ *µm* × 400 *µm* centered on the colony. A visualization of the scenario is shown in Fig. 5 (A-B).

**Fig 5.**
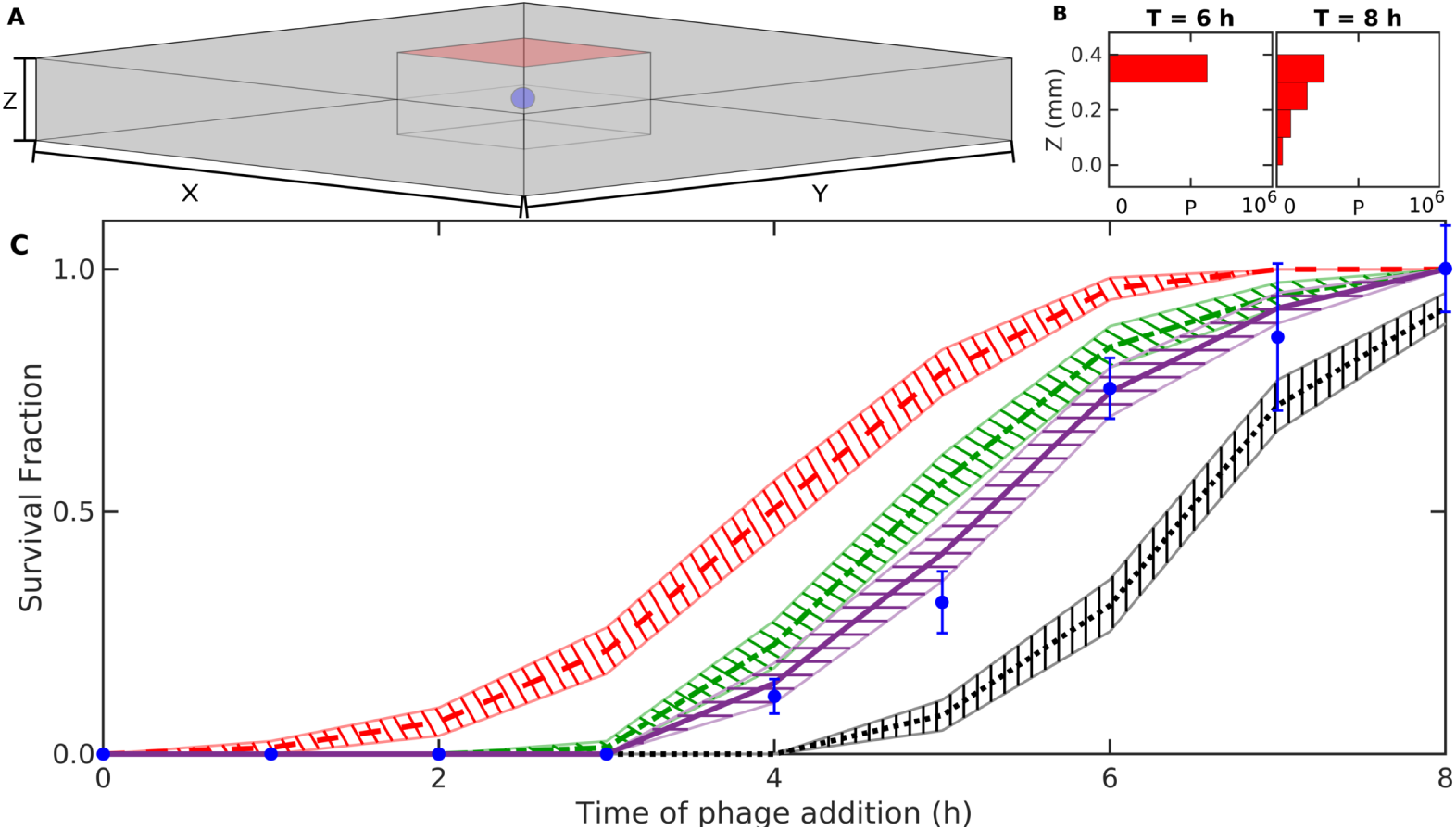
Reproducing experiment. (A) Illustration of a simulation. The spatial distribution of phages (red) at time *T* = 6.0 *h* starting from a single bacterium and with *P* = 6 · 10^5^ phages being distributed at *Z* = 400 *µm* at time *T* = 6.0 *h*. The location of the bacterial colony is shown by the blue dot. (B) The vertical distribution of phages at time *T* = 6.0 *h* and *T* = 8 *h*. (C) The fraction of colonies which grows to visible size in experiments 16 hours after phage exposure (blue error-bars) [9] and the fraction of simulated colonies (*n* = 75) which produce more than 5 · 10^5^ cells after 16 hours for *ζ* = 2.5 (red dashed line), *ζ* = 5 (green dash-dotted line), and *ζ* = 10 (purple line). As a control, we include the prediction when the colony level protections are disabled (black dotted line). The hatched area around each line indicate the standard error. The simulation here uses P1 parameters [9, 22]: *β* = 400, *δ* = 0.1 *h*^−1^, *η* = 1.32 · 10^4^ *µm*^3^/*h, τ* = 1 *h*, and parameters to match the experimental conditions [9]: Simulation volume of 400 *µm* × 10^4^ *µm* × 10^4^ *µm, D*_*P*_ = 3000 *µm*^2^/*h* (∼1/10 of phage-*λ* value) and *λ* = 60/31 *h*^−1^.

For each time, *T*_*i*_, of phage spawning, we simulate 75 stochastic developments of a single bacterial colony and compute the fraction that have grown to visible size at *T*_*i*_ + 16 hours. A colony is defined as visible, if the number of produced cells is above 5 · 10^5^, corresponding to a colony of radius ≈ 50 *µm*. Notice that we count cells produced and not the final alive cells, because dead cells also contribute to making the colony visible. Operationally the number of dead cells is counted by measuring consumed nutrient, ∫*_V_ n*(*t* = 0*h*)*dV* − ∫*_V_ n*(*t* = 16*h* + *T*_*i*_)*dV*, which has been converted to biomass.

Fig. 5 (C) compare model prediction against the experimental observations [9]. When comparing the fraction of visible colonies in Fig. 5 (C), the simulation with *ζ* = 10 reproduce the experimental result reasonably well, meaning that phages typically penetrate through 10 layers of bacteria before final adsorption.

Ref. [9] also derives a model of the critical radius, *R*_*c*_, for a bacterial colony above which it can survive a phage attack. The model assumes the colony is spherical and that the phages can penetrate a fixed distance Δ*R* into the colony. The critical radius is predicted to be: *R*_*c*_ = 3[1 + (*gτ*)^−1^]Δ*R*. From their measurement of *R*_*c*_ ≈ 25 *µm*, we can estimate Δ*R*. Since the colony is small when survival is determined, we assume the nutrient depletion is negligible and set the effective bacterial growth rate 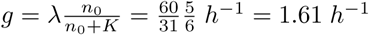. Rearranging their prediction yields the following expression:

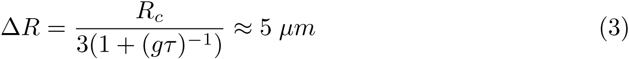

Since *Escherichia coli* has a volume of *V*_0_ ∼ 1 *µm*^3^, the parameter *ζ* gives the typical penetration depth: 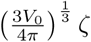, meaning we can convert the above penetration depth to a value of *ζ* = 8.3 bacterial radii.

The fraction of surviving colonies is dependent on the value of the phage diffusion constant *D*_*P*_. As we do not have an exact estimate of this parameter, we investigate how the survival fraction is affected by different diffusion constants. This is done in the appendix, where we find that if the diffusion constant is large, the fraction of surviving colonies only matches the experimental results when the shielding is strong (i.e. *ζ* small), but as the diffusion constant decreases, larger and larger values of *ζ* can reproduce the experiment.

## Discussion

The main prediction of this paper is that latency time and spatial structure greatly moderates the virulence of phages and thereby increases the robustness of phage-bacteria systems against dramatic collapses. This is in conceptual agreement with the constructed spatial study of [39] and further suggests that even the supposedly well-mixed environments of for example ref. [14] may be structured on microscopic scale. When latency time is included in the models, the phage attack is slowed and allow for bacteria survival in a relatively wide range of initial densities. Spatially varying densities improves this survivability and makes bacteria survive up to 100-fold higher phage densities, while spatial structure on the level of microcolonies allow for survival with phage densities an additional 1.000-fold higher. The geometry of the colonies reduces the phage adsorbance and shields susceptible bacteria in the center of the colony against the invading phages. In the appendix, we investigate how space alone can help bacteria survive the phage attack, and find that at low bacterial densities, space will increase survival substantially, but survival at high initial bacterial densities is governed by the latency time between phage infection and cell lysis.

Our investigation suggest that lessons learned in a well-mixed experimental system will translate poorly to scenarios where spatial heterogeneity is intrinsic. Even if the bacteria do not form microcolonies (Fig. 4(C), the bacteria will easily survive 10-fold higher phage densities. We tested scenarios where phage and bacteria are initially uniformly distributed in space, and note that if the initial distribution is more heterogeneous, the bacteria may survive even higher phage densities. One important aspect of the increased survivability of the bacteria when forming colonies, is the fundamental change to the adsorption kinetics where clustered bacteria acts as few, but larger targets. We considered the case of perfectly spherical colonies, which is the most favorable geometry for the bacteria, but clustering in general could supposedly be approximated by an adsorption term on the form: 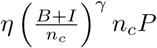, where *γ* is some volumetric scaling exponent between 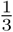 and 1.

Intuitively the fate of bacteria under a phage attack is reliant on their ability to postpone the phage attack long enough for the bacteria to consume all the available nutrient. For this reason we see a big difference when we account for the latency time *τ* (see Fig. 4(B)), since here the phage invasion is slowed, allowing more time for the bacteria to reach the carrying capacity. Spatial heterogeneity further slows the phage invasion as the diffusion of particles acts as an additional time-delay mechanism. As the consumption of the available nutrient is the only refuge for the bacteria, this suggest that a viable strategy is to grow as fast as possible, since nutrient diffuses faster than phages throughout the system. This means that even if parts of the space is completely dominated by phages, nearby bacteria might consume the common nutrient and thereby prevent the phages from fully collapsing the heavily infected areas.

Fig. 4 highlight two scenarios which lead to the elimination of bacteria. In the language of phage therapy, these situations correspond to “passive” and “active” therapy [40]. When the concentration of phages is above ∼ 10^8^ *PFU/ml* the phage load is high enough to kill the bacteria within the first few bacterial generations, corresponding to the effects of “passive” therapy, where the initial phage load eliminates the bacteria. Conversely, “active” therapy corresponds to the observed behavior of the region of the phase space which is eradicated between *T* = 0 *h* and *T* = 15 *h*. Here the phages require several replication rounds before finally eliminating the bacteria.

When the concentration of colonies becomes small (below 10^3^ *CFU/ml*), the distance between colonies become large enough that limitations of nutrient diffusion stagnates total growth (Fig. 4C,D). The nutrient on average has to travel further before meeting the colonies, which slows the overall growth. In the well-mixed cases (Fig. 4A,B) this stagnation is not seen since nutrient does not have to travel. Further, when the inoculation density is this low, the colonies would reach sizes above of 10^6^ cells and micro-gradients *within* the colony might slow growth even more than our model accounts for [8].

Our study was simplified in multiple ways. In particular, we considered the survival of one bacterial strain exposed to one phage strain. This implicitly ignores nutrient depletion due to competing bacteria, a situation which typically will favor the long term survival of any particular strain of bacteria. In addition, the parameters we have used in Fig. 2-4 does not reflect a particular phage but rather an average of the values reported for coliphages in ref. [22]. Consequently, we may have missed important dynamics in different parameter regimes. Our model also ignored the fraction of bacteria that is resistant against a given phage [41]. If there is a fraction of about 10^−5^ to 10^−6^ bacterial mutants that lack the surface receptor used by the phage, a bacterial collapse will of course not be complete.

We have found no published experiments on phage-bacteria interactions that reported cases where the susceptible bacteria are completely eliminated. In this perspective the presented modeling is a step towards a better explanation, although it still predicts conditions where bacteria should be completely eliminated. Thus further modifications to our modeling could be needed, for example by allowing the phage infections to vary hugely between the different bacteria. Perhaps the widely acclaimed variability in bacterial gene expression is a desired trait, that allow some bacteria to be resistant to phages for a substantial number of generations [42–44].

Our equations (Eq. (2)) are an approximated treatment of a system which can have quite variable strain specific properties. For example, our diverging latency time with nutrient depletion means that phage do not prey on stationary state bacteria, an assumption that is mostly correct but would fail for example for phage T7 preying on *Escherichia coli* [45]. Note that spatial structure is still relevant in such a case, as shown in protection by biofilm that hinders physical contact of phages to bacteria [25, 46]. Also the equations do not consider temperate phages, which would only induce a collapse of order 0.1 to 0.01 (typical recorded lysogeny frequencies [34]). The equations also do not consider delayed lysis, where for example the simultaneous infections by several T4 phages causes longer latency times [47]. Finally, the treatment of bacterial colonies is simplified, as we have not yet obtained a complete understanding of their protective nature [9].

A major factor in the strength of the protection mechanism is the parameter *ζ* which is a measure of the typical penetration depth of the phages. The ability of phages to penetrate bacterial colonies is likely to be dependent on several factors such as the geometry of the cells, the densities of the colonies, and whether or not the cells produce biofilm. Thereby this part of the phage protection should depend on the ability of the bacterial species to quorum sense and to collectively modify its own local environment.

The model highlights the importance of colony formation, and their ability to provide safe shelters. In fact, given that increasing colony sizes provide both local protection, and make phage percolation between colonies less likely, one should consider colony formation as a strategic option for both bacteria in the wild, our use of bacteria in industry or in human gut re-implantation, and in situations where one wants to eliminate pathogenic bacteria by phage therapy.

## Acknowledgments

This work was supported by the European Research Council under the European Union’s Seventh Framework Programme (FP/2007 2013)/ERC Grant Agreement n. 740704 and the Danish National Research Foundation (BASP: DNRF120).

