## Supplementary Material for "Sustainability of spatially distributed bacteria-phage systems"

May 13, 2019

#### Contents

**1 Adsorption to a single large target** **2**

**2 The spatial resolution** **2**

**3 The choice of shielding function** **3**

**4 The readsorption parameter** **6**

**5 The mechanisms of the colony level protection.** **7**

**6 The effect of spatial heterogeneity in the full model.** **10**

**7 The effect of latency in the full model.** **10**

**8 Phage diffusion constant and the colony survival fraction** **13**

**9 Implementation details** **14**

### 1 Adsorption to a single large target

A key component of our full spatial model, is a modification to how adsorption works on confined bacteria. When bacteria are not confined, each cell works as a small spherical sink for the phages. However, when bacteria are confined, we can consider the colony as a whole to act as a spherical sink. This difference is important, as bacteria in the center of the colony are not interacting with the phages. Mathematically, we can use a result derived by Smoluchowski[1], where he solved the rate $k$ at which diffusing particles adsorb onto a spherical sink of radius $b$:

$$k = 4\pi D b$$

If we consider the single bacterium to have radius $r_0$ then a colony containing $B + I$ members must have a volume of $V_c = \frac{4\pi}{3} r_c^3 = (B + I)\frac{4\pi}{3} r_0^3$. This means that the radii $r_c$ and $r_0$ are related by:

$$\frac{r_c}{r_0} = (B + I)^{1/3}$$

The rate of a phage adsorbing to a *bacterium* is then equal to: $\eta = 4\pi D r_0$. While the rate of a phage adsorption to a *colony* is equal to:

$$\begin{aligned} \eta_c &= 4\pi D r_c \\ &= 4\pi D (B + I)^{1/3} r_0 \\ &= \eta (B + I)^{1/3} \end{aligned}$$

### 2 The spatial resolution

Due to the diffusion constant of the nutrient in this simulation, we need to run the simulation at very small temporal resolution in order for the solution to be numerically stable. We therefore run the simulations in our paper using $\ell = 200\ \mu m$, which corresponds to a lattice of $50 \times 50 \times 50$ boxes. We test how well the simulation results has converged using this spatial resolution in Fig. S1.

We see that the relative change going from $\ell = 400\ \mu m$ to $\ell = 200\ \mu m$ is less than when going from $\ell = 1000\ \mu m$ to $\ell = 400\ \mu m$ which suggest that the simulation is converging and that the results will not change drastically when going to a smaller resolution.

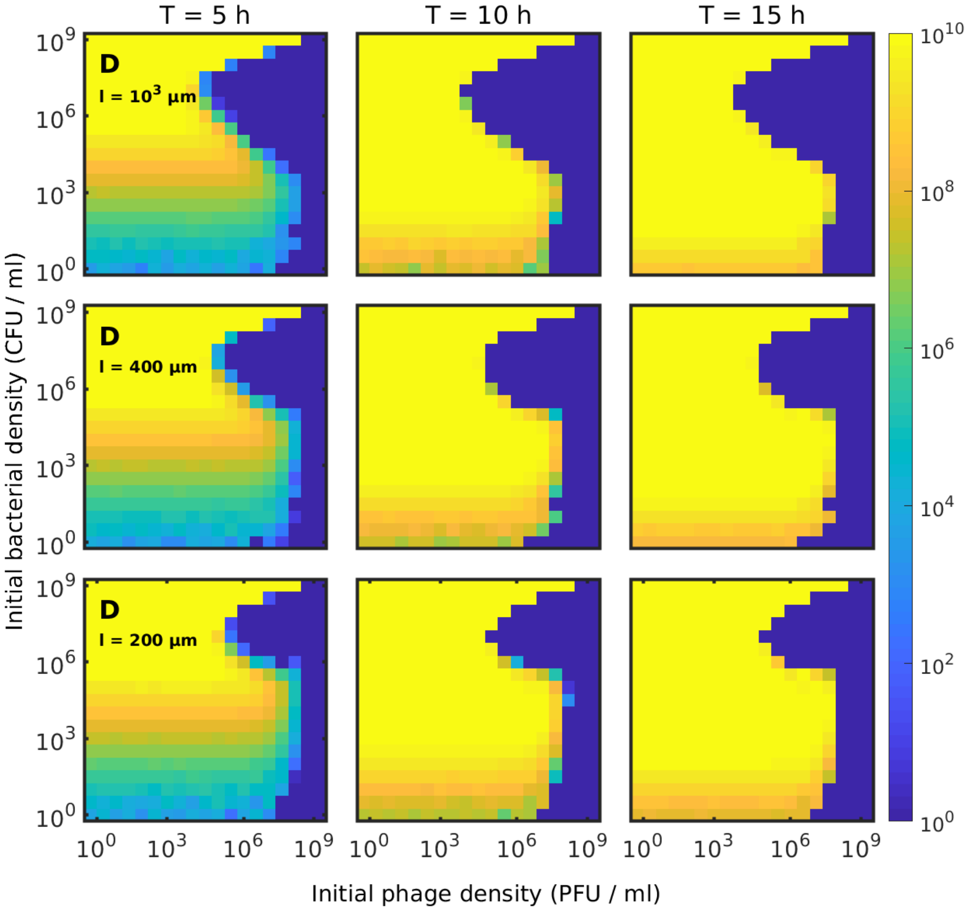


Figure S1: Simulations of the full model using different spatial resolutions: $\ell = 1000\ \mu m$, $\ell = 400\ \mu m$, and $\ell = 200\ \mu m$.

### 3 The choice of shielding function

The shielding function, $S$, represents the reduced probability of a phage hitting a non-infected cell when bacteria grows in colonies as compared to the well-mixed scenario. To build evidence for our choice of shielding function, we ran agent-based simulations of microcolonies using the framework described in ref. [2]. We have simulated micro-colonies of a various sizes ($N = \{10, 32, 100, 316, 1000\}$) and with different fractions of infected cells ($f = \{20\%, 40\%, 60\%, 80\%\}$). This allows us to compute the probability that a diffusing phage hits a non-infected cell. We then compared four different shielding functions against this data.

We first view the infected cells as small disks of area $\pi r_0^2$, where $r_0$ is the radius of a single cell. We then compare the area covered by the infected cells with the surface area of the colony: $4\pi r_c^2$. To match the small-colony limit where $n_c = B + I$ and $S = \frac{B}{B+I}$ a factor of 4 has been dropped in the expression, and we are left with the shielding function:

$$S_1(B, I, n_c) \quad = \quad 1 - \frac{I}{n_c} \cdot \left( \frac{n_c}{B+I} \right)^{\frac{2}{3}}$$

Next we again tried to compare the surface area of infected bacteria to the total surface area of the colony, but here we keep the factor of 4 when $(B + I)/n_c > 64$.

This gives our second shielding function:

$$S_2(B, I, n_c) \quad = \begin{cases} 1 - \frac{I}{n_c} \cdot \left(\frac{n_c}{B+I}\right) & (B+I)/n_c \le 64 \\ 1 - \frac{I}{4 \cdot n_c} \cdot \left(\frac{n_c}{B+I}\right)^{2/3} & (B+I)/n_c > 64 \end{cases}$$

Instead of comparing surface area, we next compare the volume of the infected cells, $I \frac{4\pi}{3} r_0^3$, with the volume of a shell of thickness $kr_0$ located on the surface of the colony: $\frac{4\pi}{3}\left(r_c^3 \left(r_c - kr_0\right)^3\right)$. This reduces to the shielding function $S_3(B, I, n_c, k)$:

$$S_3(B, I, n_c, k) \quad = \min\left(\frac{B}{B+I}, 1 - \frac{\frac{I}{n_c}}{\frac{B+I}{n_c} - \left(\left(\frac{B+I}{n_c}\right)^{1/3} - k\right)^3}\right)$$

Here we had to take the minimum of the tunneling probability and $\frac{B}{B+I}$ in order match the small-colony limit $n_c = B + I$:

Finally we created the shielding function $S_4(B, I, n_c, \zeta)$ which considers the probability of an exponentially decaying particle to move further than a distance $d = r_c - r_s$, where $r_s$ is the radius of the non-infected core. Here we also have to take the minimum to match the small colony limit:

$$S_4(B, I, n_c, \zeta) \quad = \min\left(\frac{B}{B+I}, \exp\left[-\frac{1}{\zeta}\left(\left(\frac{B+I}{n_c}\right)^{1/3} - \left(\frac{B}{n_c}\right)^{1/3}\right)\right]\right)$$

In Fig. S2, we show how each of the shielding functions match with the microcolony simulations when $\eta = 10^4\ \mu m^3/h$.

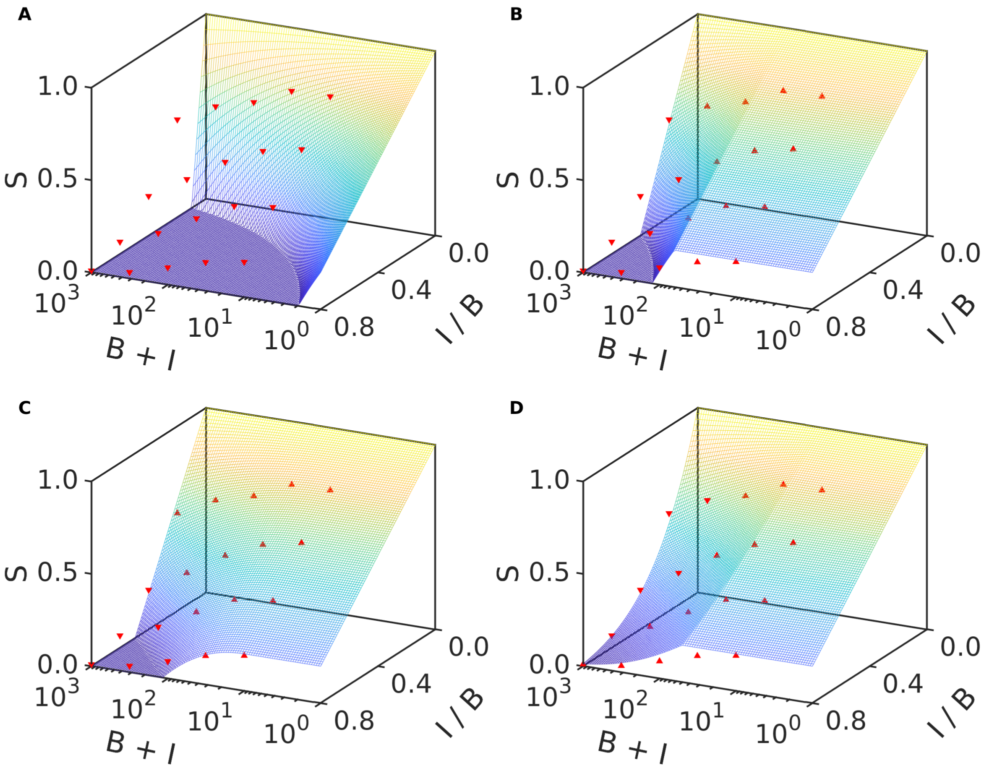


Figure S2: Comparing shielding functions with simulations. We tested four different shielding functions against simulated data for $\eta = 10^4\ \mu m^3/h$ (red triangles). For each shielding function we visualize its shape after using least-square fitting on the models with parameters ($S_3$ and $S_4$). An upwards (downwards) pointing triangle indicates that the shielding function is an overestimate (underestimate) compared with data.

Based on this analysis, we choose to use $S_4$ as the model of the shielding effect. Continuing with this model, we investigate in Fig. S3 how the parameter $\zeta$ changes as a function of the adsorption coefficient. We again use our agent-based microcolony simulations to generate the data, and fit the model to this data using least-square fitting.

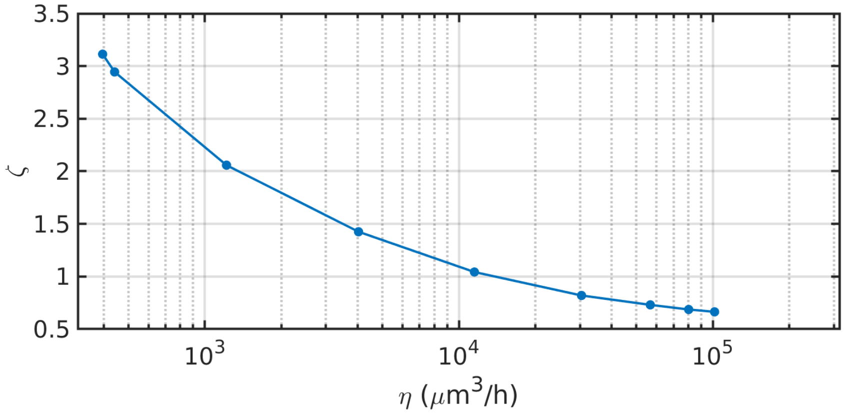


Figure S3: The absorption parameter $\zeta$ as a function of the adsorption parameter $\eta$.

The value of $\zeta$ extracted from this analysis ($\zeta \approx 1$), is not consistent with the value we find in the simulation of the experiment ($\zeta \approx 10$). This discrepancy is likely due to how the cells pack in the agent based simulations. In these simulations, we do not include any diffusion of the bacteria, meaning that they always pack tightly. In addition, the cells in the simulations are spherical instead of elongated, which might influence the packing further.

A value of $\zeta$ close to 1, corresponds with a penetration depth of one cell radius or equivalently, a penetration of $\Delta R \approx 0.6\ \mu m$. Which means it corresponds to the diffusion limited case where the phage adsorbs to the first target they meet.

In Fig. S4, we show the difference in bacterial survival between these two values of $\zeta$.

We see that when $\zeta = 5$, a large part of the phases pace goes extinct when compared to the scenario where $\zeta = 1$. The reason is that the shielding is weaker and therefore bacteria do not establish colonies of sufficient size before the phage overwhelm them.

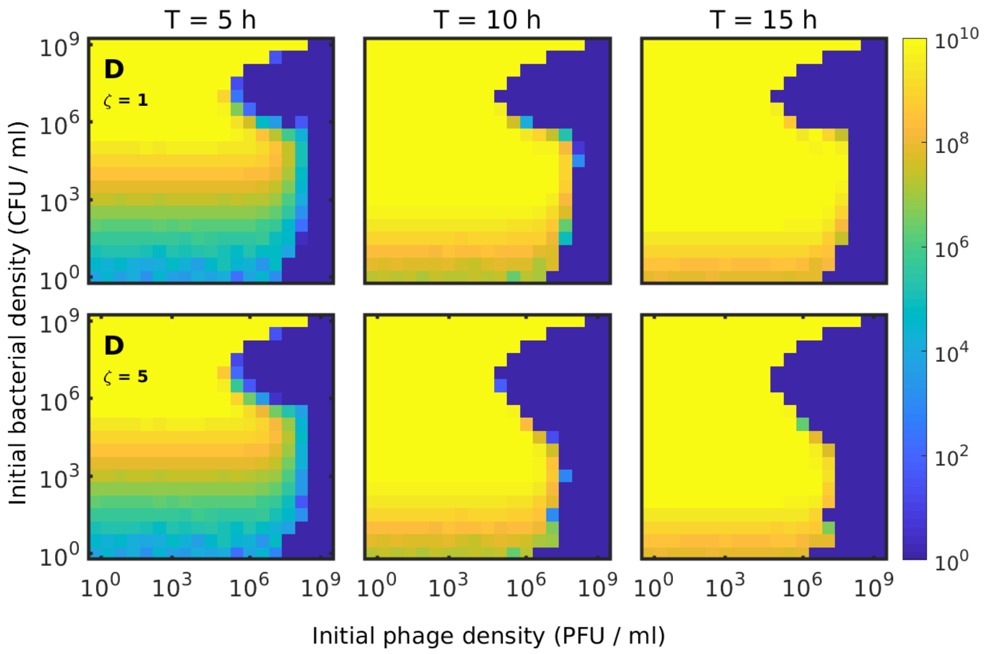


Figure S4: The influence of $\zeta$ on the ability of colonies to survive the phage attack

### 4 The readsorption parameter

In our model, the progeny phages have a probability $\alpha$ of immediately readsorbing to the colony and infect new cells. We do not have an exact estimate of what value this parameter should have, as it reflects a complicated stochastic process of how long it takes a diffusion particle to disassociate with an area of space.

We use the value $\alpha = 0.5$ as a rough estimate, and in Fig. S5 we compare with the limiting cases: $\alpha = 0$ and $\alpha \sim 1$.

When $\alpha$ is near zero, the bacterial colonies are not penalized for being close together, and are therefore very likely to survive. This scenario is clearly not realistic as any phage progeny released by lysis in a colony should lead to the new phages quickly finding the bacteria which are nearby.

As $\alpha$ approaches 1, we have the reverse scenario, where progeny phages always find the colony from which they originated. This means that the density of free phages never increases, and consequently, any colony which is not hit by the original phages are never going to be hit by any phages, which again increases the likelihood of colony survival.

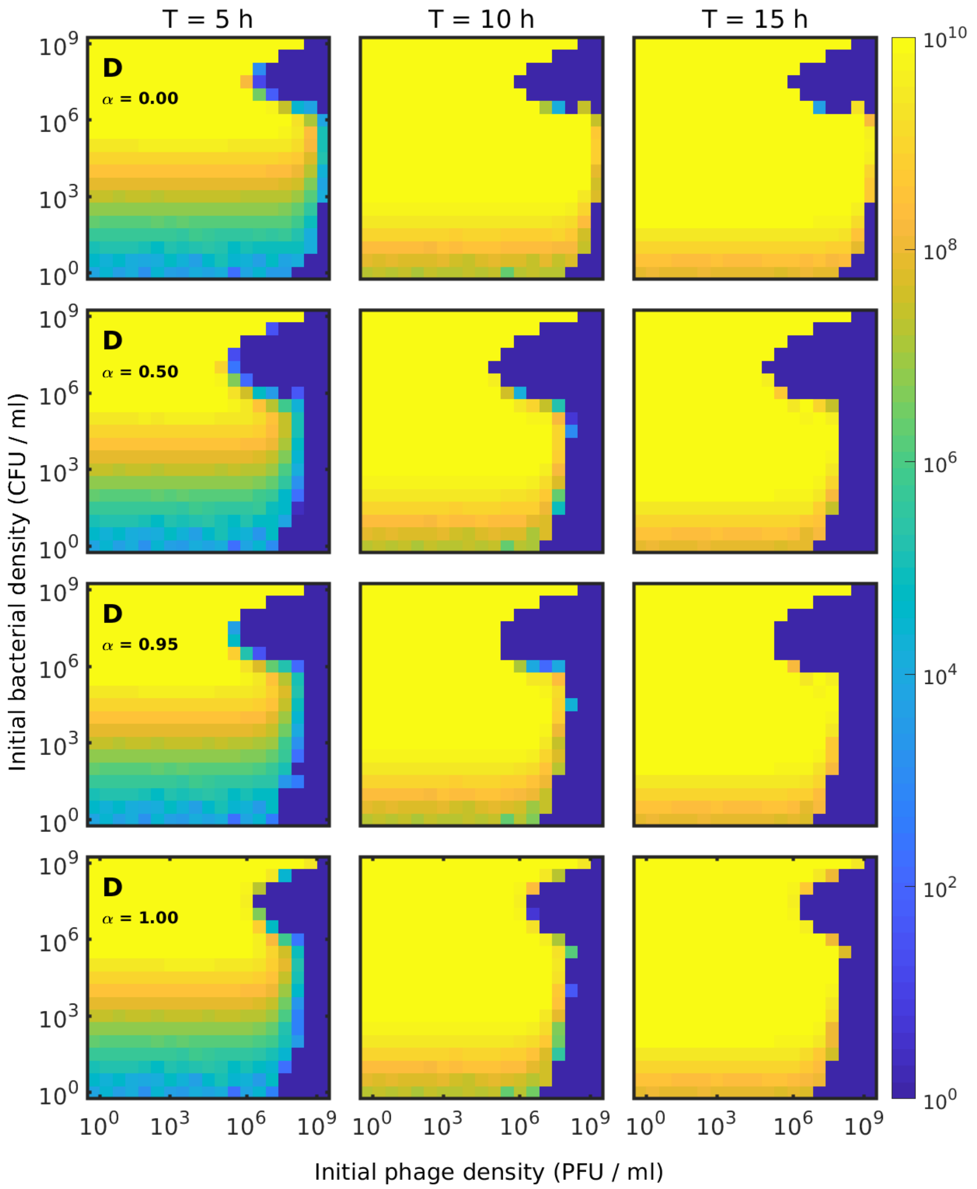


Figure S5: The effect of changing the readsorption parameter $\alpha$

### 5 The mechanisms of the colony level protection.

Going from the spatial Lotka-Volterra model in Fig. 3 (C) to the full model with colony level protection in Fig. 3 (D) added several mechanisms which influence the outcome. We here test how each of these mechanisms affect the colonies ability to survive the phage attack. The colony level protection consists of three parts:

1. Clustering
2. Shielding
3. Readsorption

Clustering was introduced as a reduction in the effective adsorption rate. I.e. we exchanged the term $\eta BP$ in Eq. (1) with the term $\eta \left(\frac{B+I}{n_c}\right)^{\frac{1}{3}} n_c$ in Eq. (2). Shielding was introduced via the function $S(B, I, n_c, \zeta)$, and readsorption is controlled by the parameter $\alpha$. Each of these mechanisms can be controlled individually, and we can test their influence one at a time. In Fig. S6, we show what each mechanism does on its own when added to the model in Fig. 3 (C).

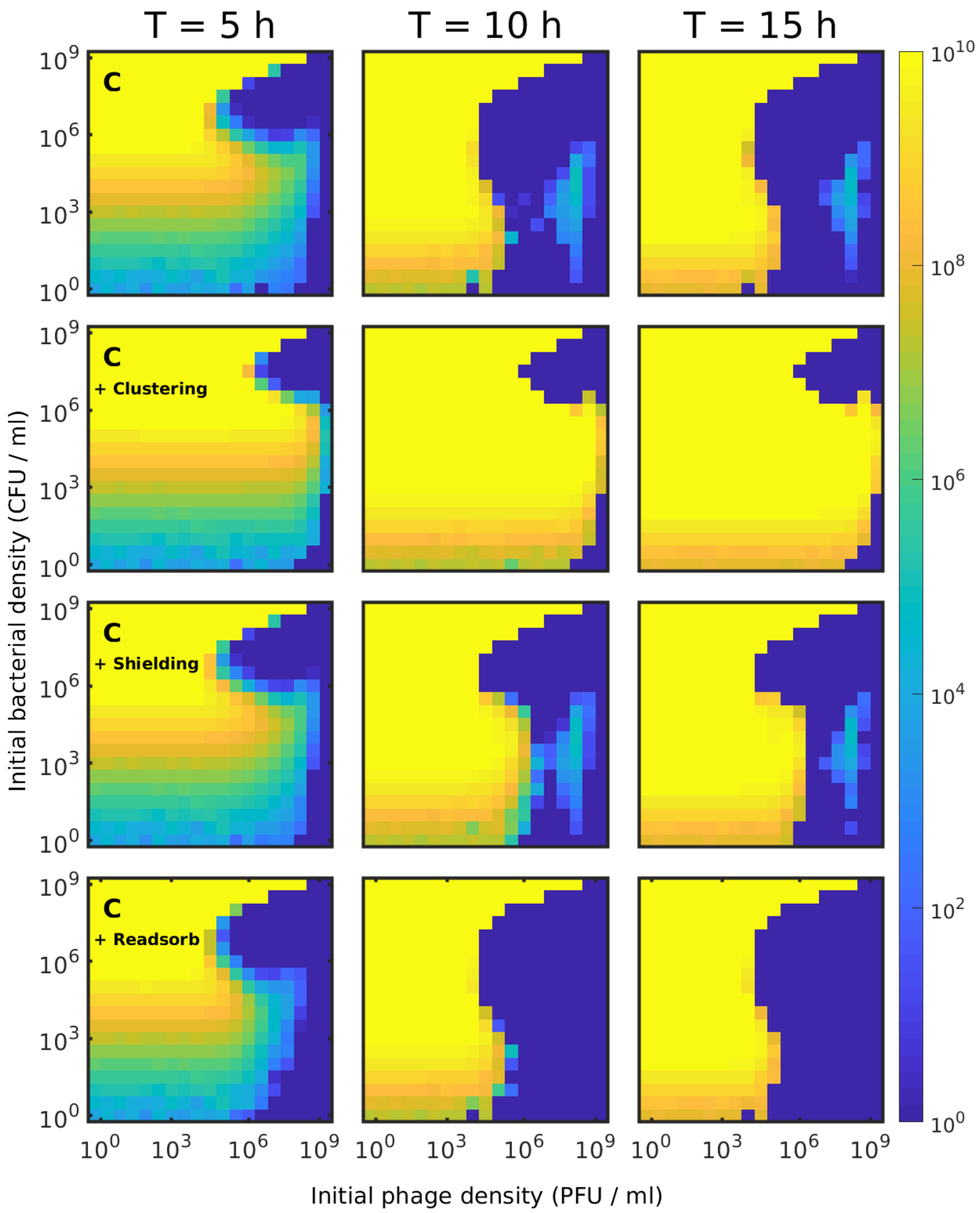


Figure S6: Introducing each effect. In this figure we test the effect each mechanism of the colony level protection (Eq. (2)).

It is clear that, when taken alone, the clustering has the biggest effect on survival, but the shielding effect also provides significant protection. Interestingly, the readsorption effect does not on its own change the survival significantly in most cases, but it does remove the marginally stable region around $P = 2 \cdot 10^8$ PFU/ml.

Next, in Fig. S7, we show how removing each of mechanism changes the survival of the full model in in Fig. 3 (D).

Here the antagonistic influence of the readsorption shows strongly. Both the shielding and clustering effects cannot on their own improve survival significantly when the readsorption is included. In both cases we see that readsorption cancels out the effect of the protection mechanism (clustering / shielding). When readsorption is removed however, the two protective mechanisms add up to yield a huge area of survival.

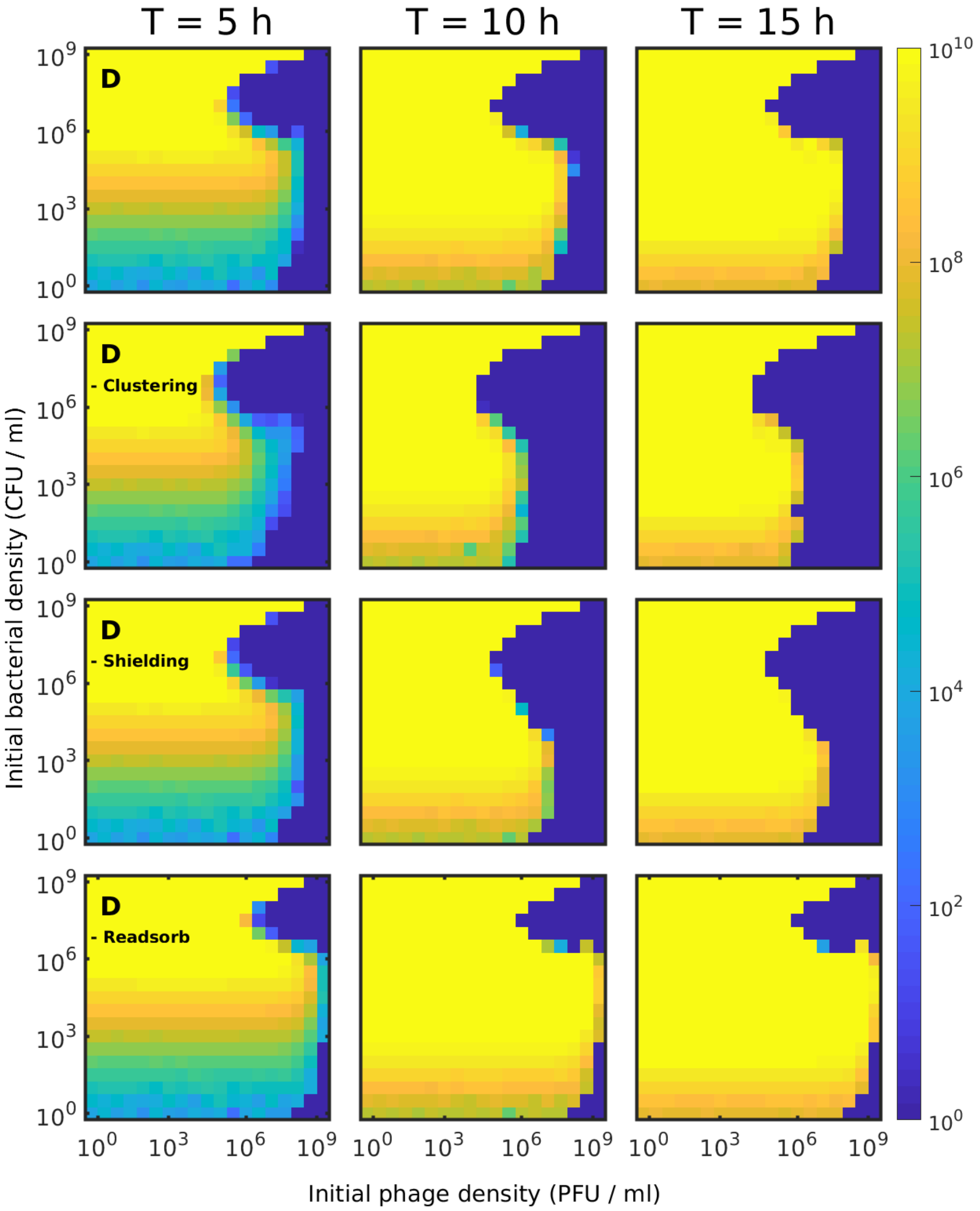


Figure S7: Removing a single effect. We test combinations of the mechanism of the colony level protection (Eq. (2)). By removing the clustering effect, we see how the shielding and readsorption effects work in unison. Next we disable shielding by setting $S(B, I, n_c, \zeta) = \frac{B}{B+I}$. And finally remove the readsorption by setting $\alpha = 0$.

#### 6 The effect of spatial heterogeneity in the full model.

The colony level protection greatly increased the survival of the bacteria, and we now test whether these mechanisms have made the effects of including spatial variation redundant. We therefore run the full model Eq. (2) within a single box of size $\ell = 1\ cm$, and compare it to the result found in Fig. 3 (D).

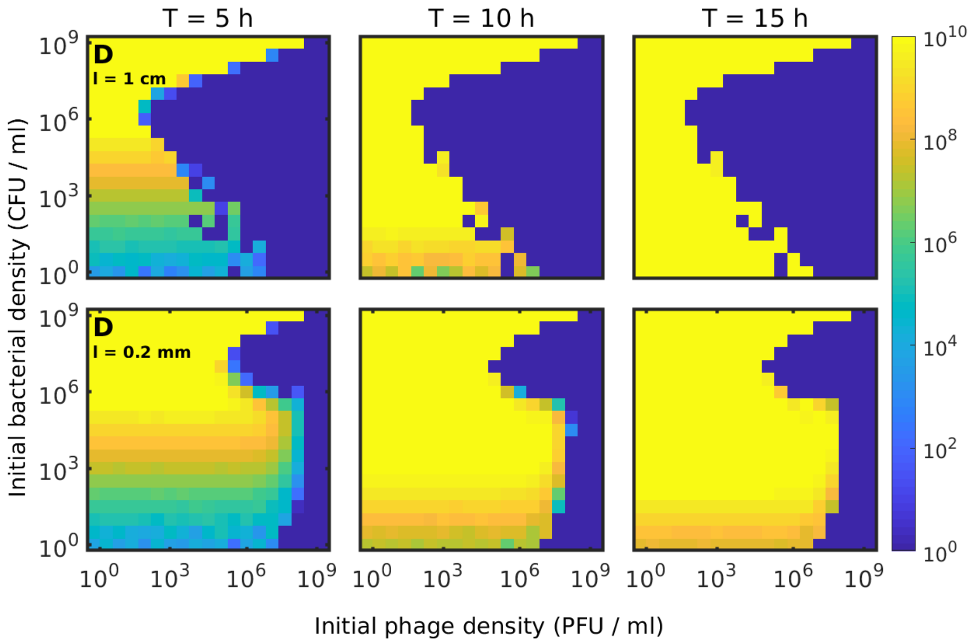


Figure S8: The effect of spatial heterogeneity on the full model. In this figure we compare Eq. (2) without the addition of spatial variation ($\ell = 1\ cm$), and with the inclusion of spatial variation ($\ell = 0.2\ mm$).

From this test, we see that the moderating effects of spatial variation is still a large factor in allowing the bacteria to survive, even with the colony level protection. One large factor of including spatial variation is to break the synchronization of events, which allows the bacterial colonies to have a distribution of sizes. Without variation, a single extremal event, such as an early adsorption or fast lysis, will amplify quickly which leads to a population collapse. The heterogeneity introduced by space is therefore needed for the protection mechanism of the model to work.

#### 7 The effect of latency in the full model.

In this paper, we are interested mostly in how the spatial effects contribute to survival of bacteria. However, we include a latency between phage infection and cell lysis in our model, since we believe this to be an important aspect of phage virulence. In this section, we test how well spatial factors alone can protect the bacteria, and we therefore simulate our model with $\tau = 0$ in the full model, and in the no-colony limit of the full model. In Fig. S9, we compare the non-spatial model with and without latency (Fig. 4A-B), with repeats of the simulations in Fig. 4C-D without latency ($\tau = 0$)

We see that spatial variation alone does provide refuge for the bacteria when the initial bacterial density is low. Here the clusters of bacteria are quite separated and the phages do not diffuse sufficiently fast to eliminate all bacteria before they consume the (bulk of the) nutrient. When including the formation of colonies, the bacteria survive at even large initial densities (of both phages and bacteria). However, in order to achieve survival at high initial densities, the latency between infection and lysis is required since it allows the bacteria to consume the nutrient before the phage invasion takes hold.

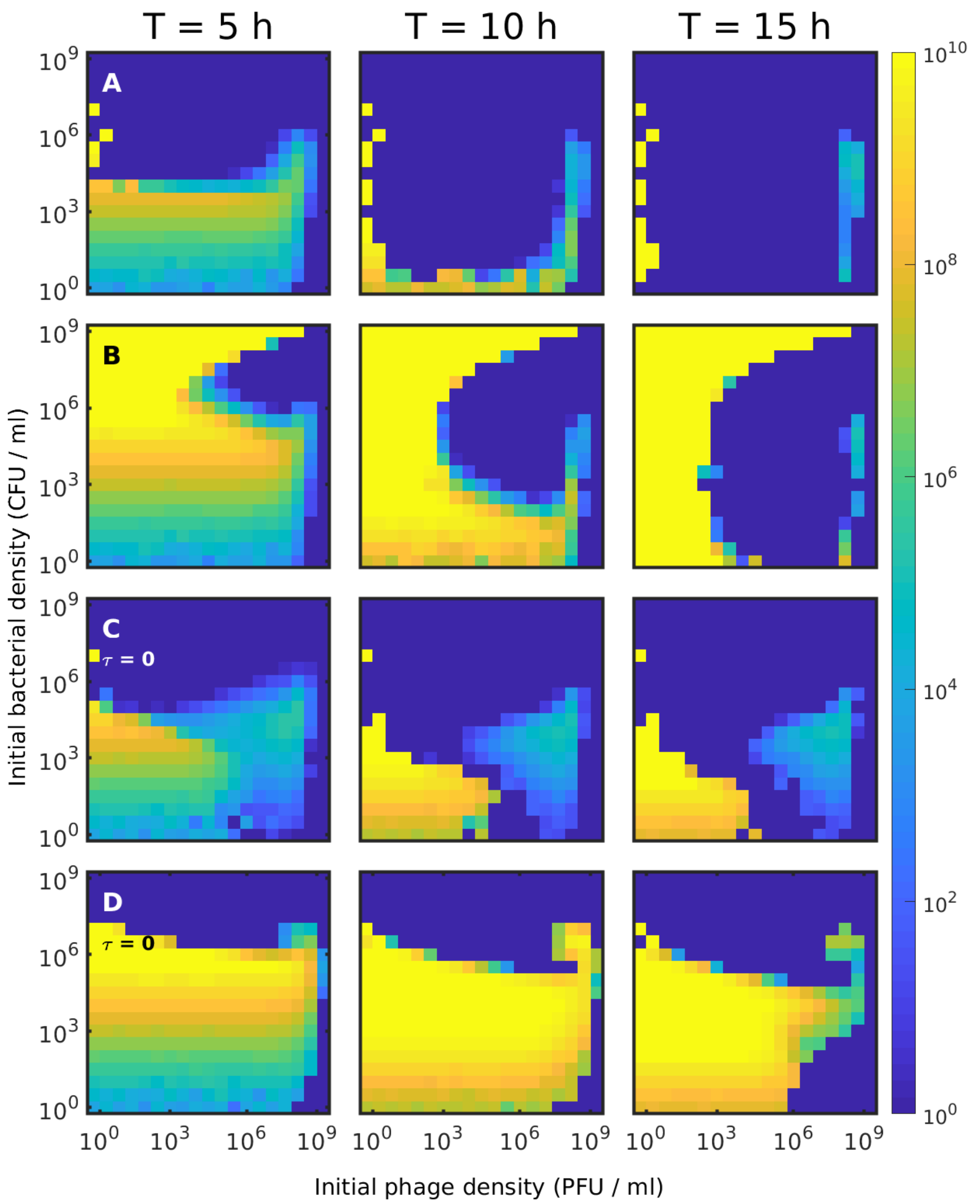


Figure S9: The effect of latency on the full model. (A) The non-spatial limiting case of Eq. (2) without latency. (B) The non-spatial limiting case of Eq. (2) with latency ($\tau = 0.5$). (C) The no-colony limiting case of Eq. (2) without latency. (D) The full model: Eq. (2) without latency.

### 8 Phage diffusion constant and the colony survival fraction

During our investigation, we learned that the diffusion constant of the phages has a large impact on the survival of the bacterial colonies. The phage diffusion constant strongly moderates the phage predation in the experimental setup we consider, since the parameter strongly effects how many phages are available to prey on the bacteria. Consequently, the value of the phage diffusion constant has an impact on which value of $\zeta$ is needed to reproduce the experimental results.

In Fig. S10, we show the fraction of surviving colonies for two phage diffusion constants ($D_P = 10000\ \mu m^2/h \sim 3\ \mu m^2/s$ and $D_P = 3000\ \mu m^2/h \sim 1\ \mu m^2/s$).

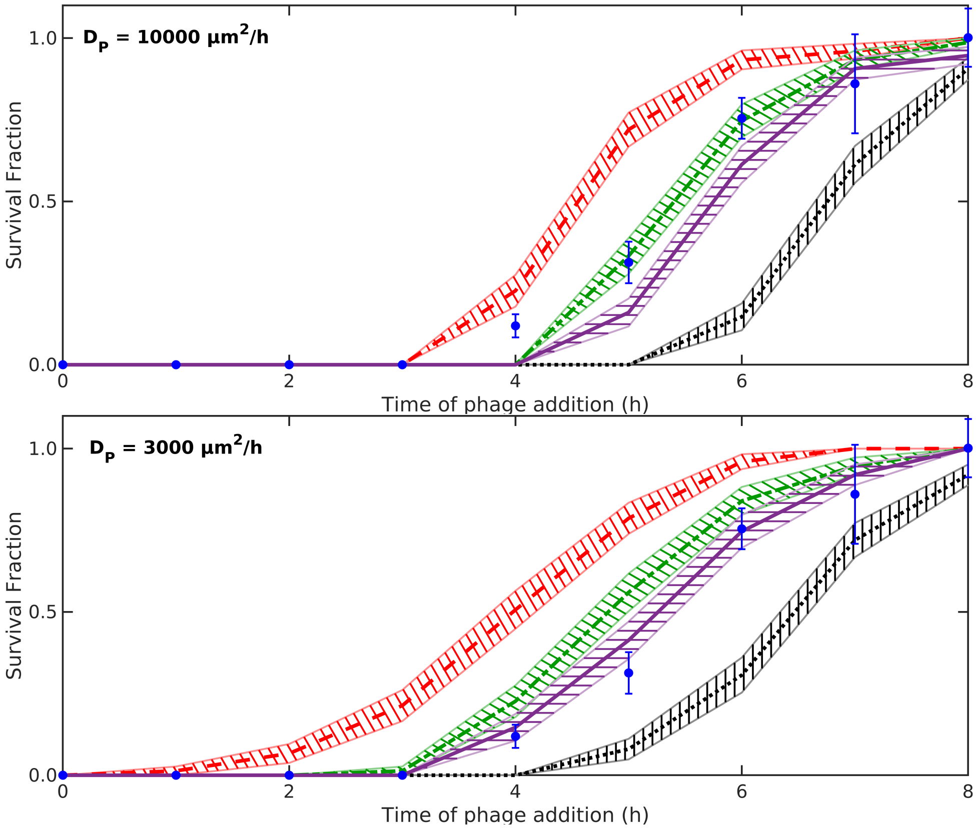


Figure S10: Phage diffusion constant and colony survival. The fraction of colonies which grows to visible size in experiments (blue error-bars) and the fraction of simulated colonies ($n = 75$) which grow to visible size for $\zeta = 2.5$ (red dashed line), $\zeta = 5$ (green dash-dotted line), $\zeta = 10$ (purple fully drawn line), and when colony level protections are disabled (black dotted line). See main text for more details.

We see that the when the diffusion constant for the phage is large, the shielding parameter $\zeta$ has to be smaller to compensate for the increased phage density in the vicinity of the colony.

### 9 Implementation details

In this section, we present the default parameters used throughout the paper (see Table S1), as well as a breakdown of the how the equations have been interpreted as describing the rates at which events occur.

If we first consider the equation which describes the dynamics of the nutrient in the system:

$$\frac{dn}{dt} = -\lambda B \frac{n}{n+K} + D_n \nabla^2 n$$

This contains two terms, a term which comes from the growth of bacteria $\lambda B \frac{n}{n+K}$ and a term which gives diffusion of nutrient $D_n \nabla^2 n$. As mentioned in the main text, the diffusion of nutrient is treated deterministically, and therefore can readily be implemented by a discrete version of the Laplace operator (forward time, central space) using a 7-point stencil. We interpret the term which describes the growth of bacteria as the rate at which new bacteria are formed. We therefore have an event (single bacterium dividing) and a rate $\rho_1$ of the event ($\rho_1 = \lambda \frac{n}{n+K}$). From the rate, we compute the probability $p_1$ that a bacterium divides in the given time-step as $p_1 = \rho_1 \cdot \Delta t$. We then draw the number of events $N_1$ which occur from a Poisson distribution with mean $p_1 \cdot B$. For each birth event that occurs, two things happen: 1) one unit of nutrient is being consumed (corresponding to the minus sign in the above equation) and 2) one new bacteria is introduced.

If we now consider the equation which gives the dynamics of the susceptible bacteria:

$$\frac{dB}{dt} = \lambda B \frac{n}{n+K} - S(B, I, n_c, \zeta) \left[ \eta \left( \frac{B+I}{n_c} \right)^{1/3} n_c P + \alpha \beta \frac{10}{\tau} I_{10} \frac{n}{n+K} \right]$$

We see the implementation of the birth event describes the first term in the equation. The remaining term is: $S(B, I, n_c, \zeta) \left[ \eta \left( \frac{B+I}{n_c} \right)^{1/3} n_c P + \alpha \beta \frac{10}{\tau} I_{10} \frac{n}{n+K} \right]$. This terms describes the number of susceptible bacteria which are being hit by phages. The phages come from two sources: free phages, $P$, and from bacteria which undergoes lysis. The term can be interpreted as a series of events, each with different rates. First we consider the event: free phages hitting the colonies. A free phage will hit a colony at a rate of $\rho_2 = \eta \left( \frac{B+I}{n_c} \right)^{1/3}$. Since we have $n_c$ colonies, the phage will hit any colony with probability $p_2 = 1 - (1 - \rho_2 \cdot \Delta t)^{n_c}$. We can get the number of phages $N_2$ which hits colonies by drawing a number from the Poisson distribution with mean $p_2 \cdot P$. Similarly, we can compute the number of bacteria which undergoes lysis, by considering the rate of which bacteria lyses: $\rho_3 = \frac{10}{\tau} \frac{n}{n+K}$. And compute the probability of lysis during the time step: $p_3 = \rho_3 \cdot \Delta t$, and then draw the number of lysing bacteria as $N_3$ from a Poisson distribution with mean $p_3 \cdot I_{10}$. For each lysis event that occurs, $\beta$ new phages are released, of which a fraction $\alpha$ readsorbs to the colonies. We can then compute the number of re-adsorbed phages which hit the colony, $N_4 = \alpha \cdot \beta \cdot N_3$.

For each of these events ($N_2 + N_4$), a phage is consumed and is removed from the simulation, and we can compute the number of phages which slip through the surface to reach the uninfected bacteria. Each phage which has hit the colony, has a probability $p_5 = S(B, I, n_c, \zeta)$ of hitting the susceptible bacteria within and we can draw the number of bacteria hit $N_5$ from the Poisson distribution with mean $p_5 \cdot (N_2 + N_4)$.

| Parameter | Value | Unit | Comment |
|---|---|---|---|
| $\Delta T$ | $2 \cdot 10^{-3}$ | h | Time increment size |
| $L$ | $10^4$ | $\mu m$ | Length of simulated space |
| $\ell$ | 200 | $\mu m$ | Length of each lattice point |
| $\lambda$ | 2 | 1/h | Maximal bacterial growth rate |
| $n_0$ | $10^9$ | 1/ml | Available nutrient (at $t = 0\ h$) |
| K | $n_0/5$ | 1/ml | Michaels-Menten constant in growth law |
| $\alpha$ | 0.5 | | Readsorption probability |
| $\beta$ | 100 | | Burst size |
| $\delta$ | 0.1 | 1/h | Phage decay rate |
| $\eta$ | $10^4/\ell^3$ | 1/h | Adsorption rate |
| $\tau$ | 0.5 | h | Lysis latency time |
| $\zeta$ | 1 | | Surface permeability |
| $D_P$ | $10^4$ | $\mu m^2/h$ | Phage diffusion constant ($\sim 3\mu m^2/s$) (T7 Diffusion rate[3]: $\sim 3\ \mu m^2/s$ and $\lambda$ diffusion rate[4]: $\sim 6\ \mu m^2/s$) |
| $D_n$ | $2.5 \cdot 10^6$ | $\mu m^2/h$ | Nutrient diffusion constant [5] |

Table S1: The default values used in the simulations.

Next, we consider the equations of the infected bacteria:

$$\frac{dI_1}{dt} = S(B, I, n_c, \zeta)\left[\eta\left(\frac{B+I}{n_c}\right)^{1/3} n_c P + \alpha\beta\frac{10}{\tau}I_{10}\frac{n}{n+K}\right] - \frac{10}{\tau}I_1\frac{n}{n+K}$$

$$\frac{dI_k}{dt} = \frac{10}{\tau}\frac{n}{n+K}\left(I_{k-1} - I_k\right) \quad i = 2, 3, ..., 10$$

At this point, we have already computed the number of susceptible bacteria hit by phages, $N_5$, and we have therefore covered the influx term in the equation of for $I_1$. The remaining terms all describe events where an infected bacterium moves from one stage to the next stage. All of these events have the same rate as we have seen from $I_{10}$, namely the rate $\rho_3$. We can then compute the number of bacteria which moves out from state $I_k$ as $N_{k+6}$ drawn from a Poisson distribution with mean $p_3 \cdot I_k$.

Finally, we can consider the equation that describe the phages:

$$\frac{dP}{dt} = (1-\alpha)\beta\frac{10}{\tau}I_{10}\frac{n}{n+K} - \delta P - \eta\left(\frac{B+I}{n_c}\right)^{1/3} n_c P + D_p\nabla^2 P$$

Here we have computed the number of bacteria lysing $N_3$, and the first term in the equation has therefore already been covered. The second term $-\delta P$, describes the number of phages which decay over time. Here we can compute the number of phages which decay $N_{16}$ from the Poisson distribution with mean $\delta \cdot \Delta t \cdot P$. The third term has been covered by $N_2$ when we computed the number phages which hit the colonies. Finally we have the diffusion term $D_p\nabla^2 P$, which treat as random walk where each phage has a probability $2\frac{D_P\Delta t}{\ell^2}$ of jumping the a neighbour point. By drawing from a Poisson distribution, we can compute the flux, of phages in to / out of the grid point.
